# Evaluating the Role of Cell-free Fetal DNA in Inflammation and Spontaneous Preterm Birth

**DOI:** 10.1101/191528

**Authors:** Sara R van Boeckel, Heather MacPherson, Donald J Davidson, Jane E Norman, Sarah J Stock

## Abstract

Preterm birth is the leading cause of neonatal mortality. While spontaneous preterm birth (sPTB) is the cause of over 70% of PTB, the pathogenesis behind sPTB remains unclear. Cell-free fetal DNA (cff-DNA) originates from the placenta and is increased in women who develop PTB. It has been demonstrated that fetal DNA is hypomethylated and is pro-inflammatory. The pro-inflammatory properties of placental-derived DNA, the effects of placental inflammation on the production of cff-DNA, and its significance in the pathogenesis of PTB are unknown.

Using a human placental explant model, we analysed the effect of lipopolysaccharide (LPS) stimulation on cff-DNA production, and used the cff-DNA generated by these explants to examine the methylation profile and *in-vitro* pro-inflammatory properties of cff-DNA. LPS caused significant production of TNF-α from placental explants, but did not significantly increase the cff-DNA production. Placental-derived cff-DNA, was found to have a small proportion of unmethylated CpG motifs, but was more similar to adult DNA than to more highly unmethylated E-coli DNA. However, cff-DNA did not elicit production of inflammatory cytokines (IL-6, IL-8, TNF-α and CXCL10) by peripheral blood mononuclear cells from pregnant women. Furthermore, in contrast to LPS, intra-uterine injections of mouse placental DNA did not decrease time to delivery in an *in-vivo* mouse PTB model compared to control animals.

This study demonstrates that placental inflammation does not increase the production of cff-DNA in placental explants, and cff-DNA alone is not sufficient to elicit an inflammatory response in human PBMC cultures *ex-vivo.* It also shows that mouse placental DNA does not cause PTB *in-vivo.* This suggests that cff-DNA might be predominantly an effect of parturition and not a principal causative agent.

## Introduction

Preterm birth (PTB), defined as delivery before 37 weeks of gestation, is the leading cause of neonatal morbidity and mortality (1). There are currently few effective therapies to prevent preterm birth (1, 2). It is clear that inflammation plays a central role in the induction of spontaneous PTB (sPTB), which contributes to over 70% of total PTBs (3, 4), but the key events in the initiation and regulation of this inflammation are unknown. Recent studies have suggested that high levels of circulating cell-free fetal DNA (cff-DNA) may be a trigger for inflammation, and consequent inflammation-induced parturition, through activation of toll-like receptor (TLR)9 (5-7).

Cff-DNA is of placental origin, generated during cell-death of the trophoblasts in the syncytiotrophoblast layer (8-11). The levels of circulating cff-DNA increase throughout gestation (12, 13), and higher levels are associated with various pregnancy pathologies; including pre-eclampsia and growth restriction, when compared to normal pregnancies (9, 14, 15). Studies have also shown higher levels of cff-DNA in women who deliver preterm compared to women who deliver term (16-18). *In-vitro* studies have demonstrated that inflammation, induced by lipopolysaccharide (LPS), can increase programmed cell-death (apoptosis) in placental explants (19), and can increase cff-DNA release in mouse placental explants (20).

Cff-DNA differs in composition to circulating cell-free DNA of maternal origin, being less methylated than adult DNA (6, 7). Because the innate pattern recognition receptor TLR9 is activated by unmethylated CpG oligonucleotide (ODN) sequences (21), this makes cff-DNA a potential TLR9 ligand. Cell-free DNA of tumor origin has been shown to cause inflammatory responses in epithelial cells through TLR9 and the Stimulator of Inteferon Genes (STING) pathways (22, 23), however, it remains unclear whether circulating cff-DNA is indeed pro-inflammatory *in-vivo* in pregnancy.

We hypothesized that *in utero* inflammation would increase cff-DNA release from the placenta into the circulation, where it could activate the maternal immune system and contribute to the induction of sPTB. The aim of this study was to examine the effect of LPS on cff-DNA release in a human placental explant model, investigate the pro-inflammatory effects of cff-DNA on maternal peripheral mononuclear cells, and determine whether placental DNA an induce PTB in a mouse model.

## Materials and Methods

### Human samples

Placental tissues and whole blood from pregnant women were collected under Edinburgh Reproductive Tissue Bio Bank ethical approval (ERTBB 13/ES/0126). Samples were donated by women with term pregnancies (37-40 weeks gestation) who were not in labour, had a BMI < 30, with no pregnancy complications, no blood borne diseases, no immunosuppressive disease or medication, and no current signs of infection.

### Placental Explant Culture and Stimulation

Within 30 minutes of delivery, 10 x 10 cm of placental tissue with no visible areas of calcification or hemorrhage was dissected and immediately immersed in cold Dulbecco’s phosphate-buffered saline (DPBS; Gibco). Two washes were performed with DPBS, followed by a wash in culture medium (RPMI (Gibco), 10% Fetal Bovine Serum, 1% penicillin/ streptomycin). While immersed in culture medium, placental pieces of approximately 150 mg were separated, using forceps and knife, into separate 6-well plates (one villous explant of 150 mg per well) and then cultured at 37°C, 5% CO_2_. These placental explants were then stimulated with LPS (derived from E-Coli (0111:B4 Sigma) at 2 ng/ml, 20 ng/ml or 200 ng/ml for 1, 6 and 24 hours. Supernatants were collected for cff-DNA extraction and quantification.

### Cell-free fetal DNA extraction and quantification

Supernatant was harvested from placental explant cultures and cff-DNA extracted using the QIAamp Circulating Nucleic Acid Kit (Qiagen). The manufacturer’s protocol was followed, using 1 μg carrier RNA and eluting in buffer AE (10 mM Tris-Cl, 0.5 mM EDTA; pH 9.0). Quantification of the cff-DNA isolated was performed with an absolute quantitation Real-Time multiplex PCR machine (ACB/SRY), over seven serial dilutions, using a standard curve of placental DNA. Cff-DNA was quantified in pg/μl per mg of villous tissue in each placental explant culture.

### Placental and Adult DNA extraction

Human placental DNA was extracted from placental tissue, which had been snap frozen at time of collection, using the Qiagen Blood & Tissue DNA extraction kit with prolonged lysing period for increased fragmentation (2 x 2 min at 25Hz). This DNA was used as a control for cff-DNA validation and quantification, and as a control stimulus in parallel to cff-DNA in *in-vitro* cell stimulation experiments. Mouse placental DNA was extracted from mouse placentas at gestational ages 17-19 days using the same method as human placental DNA. Mouse placental DNA was concentrated by precipitation with 3 M Sodium Acetate pH 5.2 and 100% ice-cold ethanol and eluted in sterile DPBS. Adult DNA was extracted from whole blood from a healthy non-pregnant woman (ERTBB 13/ES/0126) in sodium citrate tubes. The kit described above was used following manufactures protocol for whole blood DNA extraction. Adult DNA, Human and mouse placental DNA were quantified using Nano-spectometry.

### Peripheral Blood Mononuclear Cells (PBMC) isolation and stimulation

Whole blood was acquired in sodium citrate tubes. PBMCs were harvested using Histopaque with a density gradient of 1.07 g/ml, washed twice with PBS, and finally washed with the culture medium (RPMI with 10% FCS and 5% Penicillin/Streptomycin). Cell number and viability was assessed using a haemocytometer and Trypan blue staining. PBMCs were plated at 2.0 x10^6^/ml in 12-well plates and maintained at 37°C, 5% CO_2_ for three hours, before stimulation with 1 μg/ml CpG oligodeoxynucleotides 2395 (ODN; Invivogen), 1 μg/ml E-coli DNA (dsDNA E-coli Invivogen), or 0.5 μg/ml cff-DNA. Supernatants were collected after 18 hours culture at 37°C, 5% CO_2_.

### Enzyme-Linked Immunosorbent Assay (ELISA)

R’D Duoset ELISA’s for IL-6, IL-8, TNF-α, CXCL-10 were used for cytokine analysis, according to the manufacturer’s protocols. Supernatants from PBMC stimulation studies were used after being centrifuged for 1 minute at 9400 x*g* to spin down any residual cell debris.

### Methylation quantification

Whole DNA methylation quantification was performed using Epigentek MethylFlash Global DNA Methylation (5-mC) ELISA Easy Kit (Colorimetric). The manufacturer’s protocol was followed. Whole DNA was methylated using M.SssI methyltransferase (New England BioLabs), using manufacturer’s protocol, with methylation halted by incubation at 65°C for 20 minutes. This yielded the maximum amount of methylated CpGs. The level of unmethylated CpGs in starting materials was quantified by subtracting the amount of methylation quantified without use of the methylation enzyme from the amount of methylation quantified after use of methylation enzyme (maximum methylation).

### In-vivo experiments

*In-vivo* studies were conducted under a UK Home Office license to J.E.N. (60/4241) in accordance with the Animals Scientific Procedures Act (1986). C57Bl/6 virgin female mice were purchased from Charles River Laboratories (Margate, UK). Mice were housed in grouped cages of 6 mice per cage under normal husbandry conditions and acclimatised for a minimum of 1 week before use. Female mice (aged 6-7 weeks) were mated with C57Bl/6 stud males. The presence of a vaginal copulatory plug indicated day 1 of gestation. Pregnant mice with no appearance of a plug were not used in experiments, as gestational age was uncertain.

An intra-uterine ultrasound model previously optimised in our group was used (32). Briefly, mice underwent anaesthesia with isoflurane inhalant for the duration of the procedure (5% for induction, 1.5% for maintenance in oxygen). After removing abdominal hair with depilatory cream, warmed ultrasound gel was applied. Scans were performed with the Vevo 770 high-frequency ultrasound scanner (FUJIFILM VisualSonics, Inc., Toronto, ON, Canada) with a RMV 707B probe (center frequency, 30 MHz). At gestational age day 17, space between two intra-amniotic sacs was located with ultrasound where ultrasound-guided injection was administered; delivering 25 μl of sterile DPBS, or mouse placental DNA (3 μg/dam, 30 μg/dam or 300 μg/dam in 25 μl in DPBS). In a separate cohort, mice were injected with 1 μg/dam LPS (0111:B4 Sigma) in 25 μl DPBS. Mice were monitored to recover on warm mats in an individual cage. After recovery, mice were monitored using closed circuit television cameras and a video recorder. Time to delivery was measured as the time between injection and delivery of first pup. Mice were individually housed and therefore assessed as the experimental unit in statistical analyses.

### Statistics

Data in graphs are show as means ± SD. *In-vitro* and *in-vivo* stimulations were analysed using one-way or two-way ANOVA with Dunnet’s or Tukey’s multiple analysis post-test. Statistical analyses were performed with GraphPad Prism version 7.0 (GraphPad, San Diego, CA). P < 0.05 was considered statistically significant.

## Results

### LPS-induced inflammation does not increase cff-DNA release from human placental explants

Placental explants, cultured with 2, 20 or 200 μg/ml of LPS or vehicle (no stimulation) for 1, 6 or 24 hours, showed a significant and dose-dependent induction of TNF-α release, confirming the activity of LPS (figure 1A). Placental explants showed a significant increase in cff-DNA release at 24 hours compared to the 1-hour samples (N = 3), but this was observed to be irrespective of LPS stimulation (figure 1B). In order to more closely evaluate the effect of LPS at a time point prior to high spontaneous cff-DNA generation, dose response experiments were repeated at 6-hours (N = 6, figure 1C). No significant effect of cff-DNA release was seen at any of the concentrations of LPS used. These data suggested that LPS-mediated inflammation is not sufficient to induce the release of cff-DNA in placental explant cultures.

**Figure 1:**
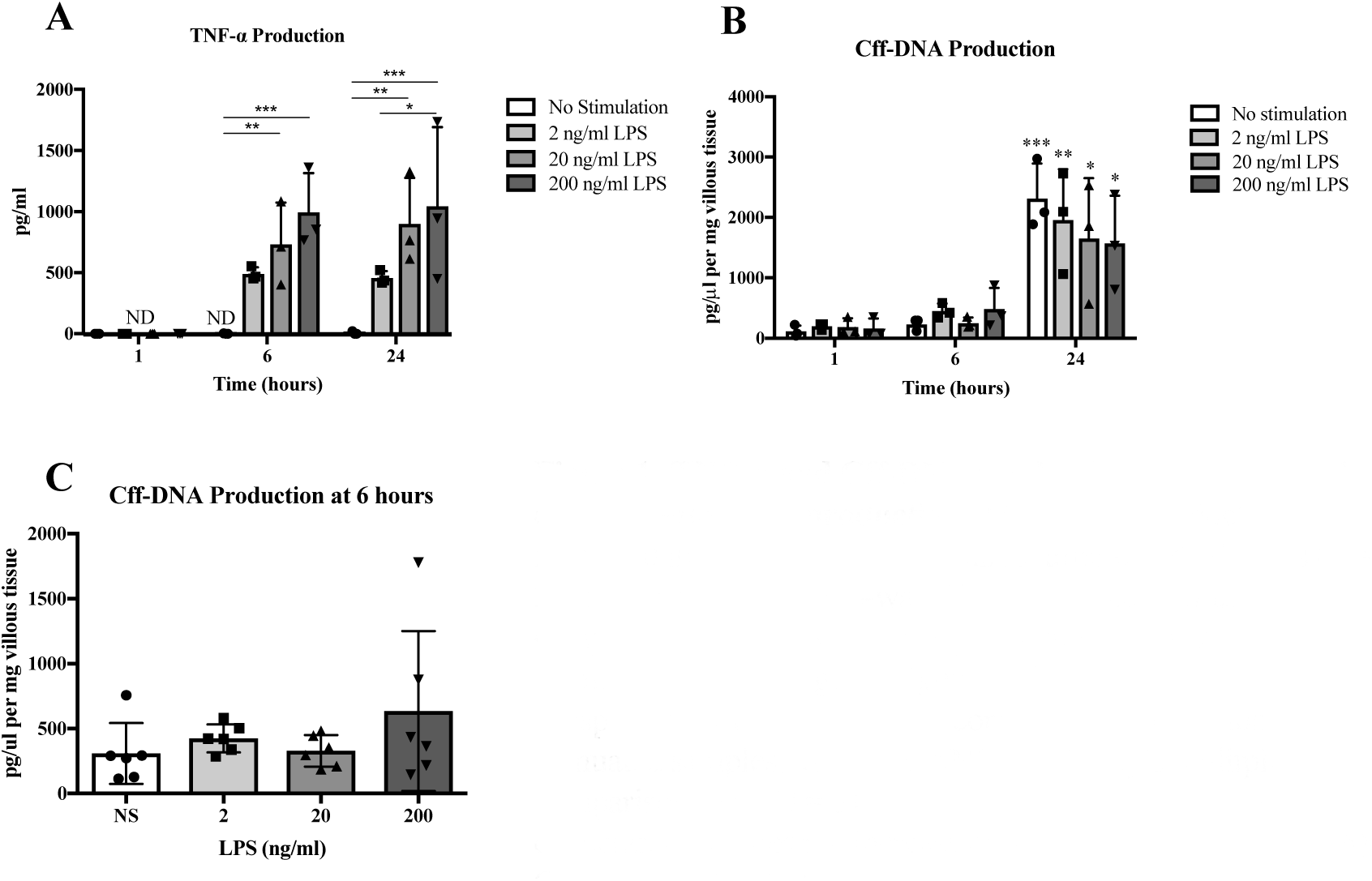
TNF-⁣ and Cff-DNA production from placental explants after LPS production. (A) TNF-⁣ production increased in a dose-dependent manner after 6 and 24 hours (N= 3) exposure to LPS. Two-way ANOVA, Tukey’s multiple comparisons test (**B)** Cff-DNA production (measured by quantifying the SRY gene) significantly increased at 24 hours irrespective of LPS stimulation compared to 1 hour non-stimuated samples, two-way ANOVA, Tukey’s mulitple comparisons test (N = 3). (**C)** LPS stimulation at different concentrations does not increase cff-DNA after 6 hours (N = 6) One-way ANOVA. All bars represent mean with ±SD, * = p ⁣ 0.05; ** = p ⁣0.01, *** = p ⁣.001.

### Cff-DNA does not cause inflammation in human PBMCs from pregnant women

To assess the ability of cff-DNA to elicit an inflammatory response, PBMCs isolated from pregnant donors were cultured for 18 hours in the presence of 500 ng/ml cff-DNA, generated *ex-vivo* from unstimulated cultured placental explants after 24 hours of culture. CpG ODN (ODN; 1 μg/ml), E-coli DNA (1 μg/ml) or LPS (500 ng/ml) were used as positive controls. PBMCs stimulated with cff-DNA showed no difference in the levels of IL-6, CXCL10, TNF-α, or IL-8 produced (N = 3-14), compared to vehicle-treated controls. In contrast PBMCs stimulated with LPS showed a significant increase in IL-6, TNF-α and IL-8 production, and ODN induced significant CXCL10 production (figure 2). These *in vitro* data do not support the hypothesis that cff-DNA would induce systemic inflammation in pregnancy.

**Figure 2:**
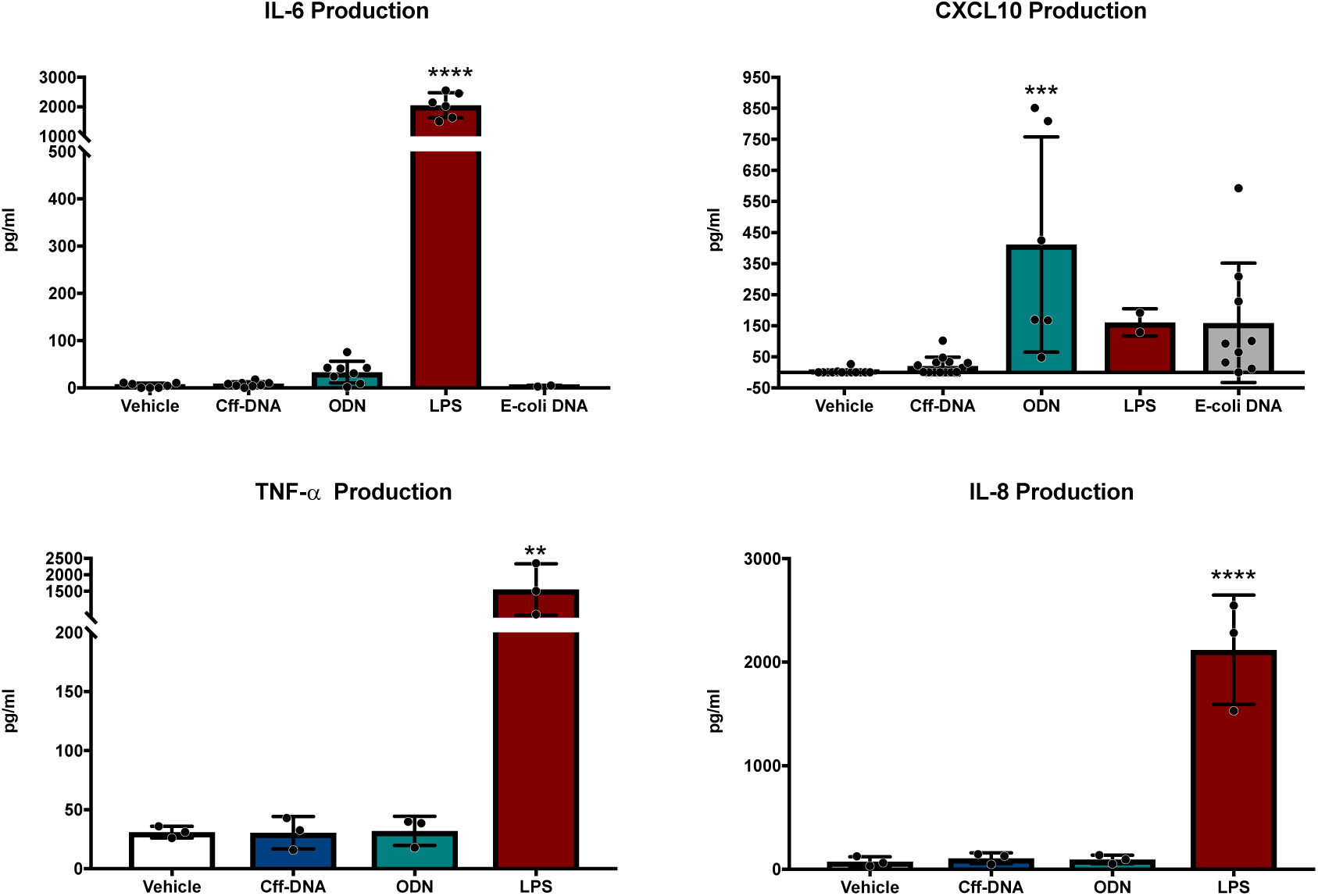
Cytokine and chemokine production by Peripheral Blood Mononuclear Cells from pregnant women after 18 hours. Data represent means ± SD. One-way ANOVA with Dunnett’s multiple comparison test was performed for all cytokines and chemokines; **, p ≤ 0.01, ***, p ≤ 0.001, ****, p ≤ 0.0001 compared to vehicle. Cff-DNA = cell-free fetal DNA derived from placental explant method (500 ng/ml), LPS = lipopolysaccharide (500 ng/ml), ODN = TLR9 agonist CpG oligonucleotide (1 μg/ml), E-coli DNA (1 μg/ml).

### Methylation of Cff-DNA

In order to determine the methylation status of the cff-DNA used in our studies, we quantified the proportion of CpG islands that were hypomethylated on the cff-DNA, placental DNA, and adult human DNA, and compared this with E-coli DNA (table 1). These analyses demonstrated that ~80% of the CpG motifs on E-coli DNA (a known TLR9 ligand) were unmethylated; with 1.9% of the total DNA comprised of methylated CpG islands, with a further 9.1% unmethylated CpG. In contrast for cff-DNA, placental and adult DNA, there was minimal evidence of unmethylated CpG islands (although 0.06% quantified for cff-DNA), but 1.5% of the total DNA was comprised of methylated CpG islands. These data confirm that E-coli DNA is heavily unmethylated compared to human DNA, but find no evidence that cff-DNA is less methylated than adult DNA.

**Table 1:** Quantification of total DNA methylation and unmethylated CpGs. Results shown as percentage of total DNA. Cff-DNA = cell-free Fetal DNA.

| DNA Sample | % Methylated CpG Islands | % Unmethylated CpG islands |
| --- | --- | --- |
| Adult DNA | 0.34 | 0 |
| Cff-DNA | 1.44 | 0.06 |
| Placental DNA | 0.82 | 0 |
| E-coli DNA | 1.93 | 9.10 |

### Mouse Placental DNA does not cause sPTB in an intra-uterine PTB model

To assess the effect of placental DNA on time to delivery, we administered different concentrations of mouse placental DNA into the uterus of pregnant mice (at day 17 gestation) by ultrasound guided injection (N = 11) and compared the time to delivery with control injections of saline (N = 4) or LPS (1 μg/dam) as a positive control (N = 4). There was no difference in the time to delivery in mice administered intrauterine placental DNA versus saline controls (58.6 ± 3.5 hours for saline, versus 59.0 ± 7.1 in 3 μg/dam, 54.8 ± 11.1 in 30 μg/dam and 67.4 ± 14.3 in 300 μg/dam; figure 3). No placental-DNA or control treated mice delivered preterm (defined as delivery within 36 hours of injection). In contrast, mice treated with 1 μg/dam of LPS had significantly reduced time to delivery (mean time to delivery = 19 hours; figure 3).

**Figure 3:**
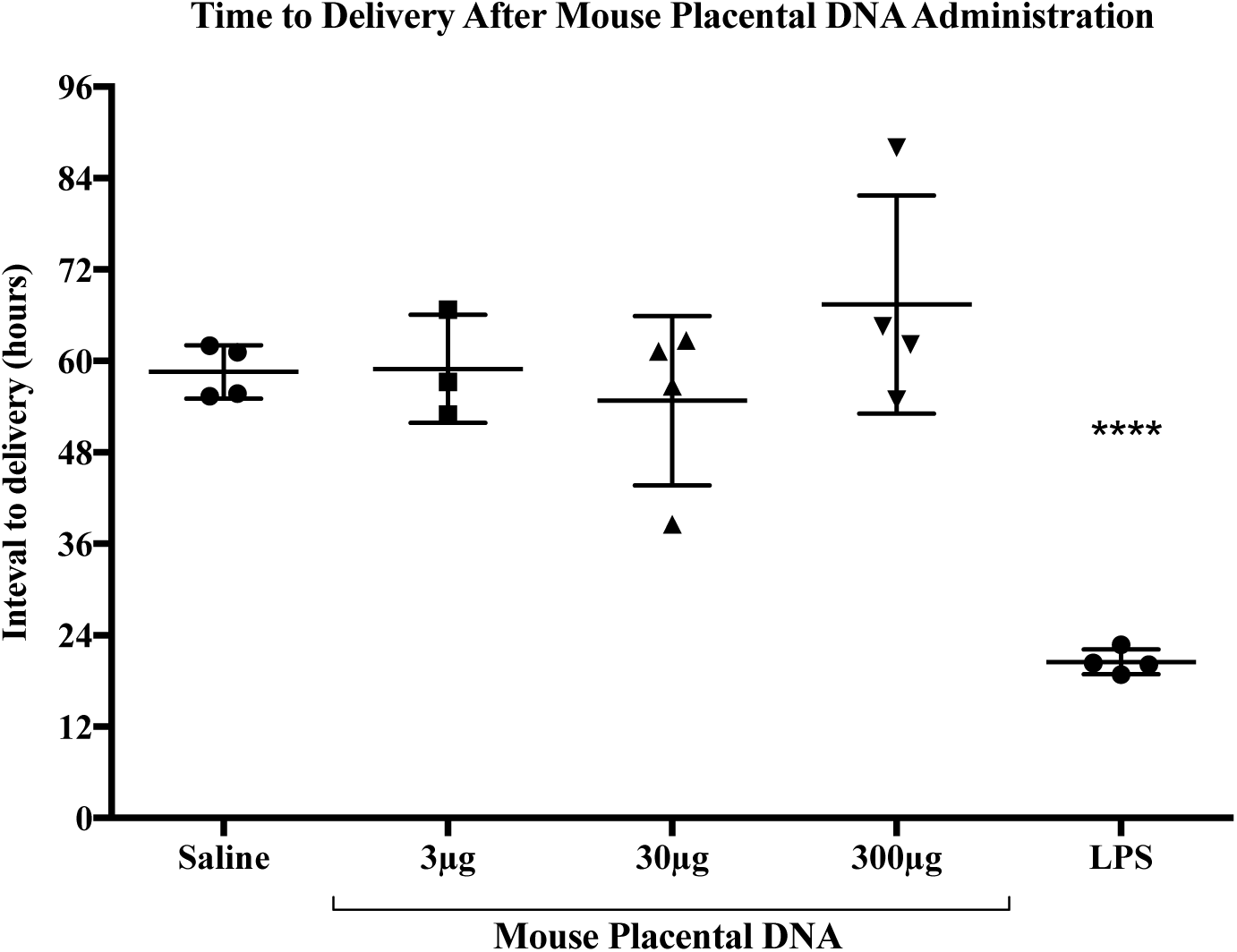
Interval to delivery after intrauterine injection of mouse placental DNA. On day 17 of gestation mice were injected with either saline. LPS. or mouse placental DNA in 25 μ saline. Substances were injected between two amniotic sacs, identified through ultrasound. Time to delivery was acquired using CCTV cameras. Mean and standard deviations are shown. One-way AXOVA was performed followed by Dunnet’s multiple comparison test compared to saline *** = p < 0.0001.

## Discussion

This study demonstrates that cff-DNA from human placental origin is not proinflammatory to PBMCs from pregnant women *ex-vivo*, and that murine placental DNA does not cause sPTB in a mouse model. These findings suggest that cff-DNA alone is not sufficient to elicit an inflammatory response that can lead to sPTB.

To our knowledge, this is the first study to use placental-derived cff-DNA from supernatants of placental explants in combination with a specific cell-free DNA extraction kit in *in-vitro* models. The cff-DNA acquired had been released by placental explants and was therefore thought to be more comparable to that observed in systemic circulation during pregnancy. Human fetal DNA (7) and mouse placental DNA (6), both extracted from tissue, have been shown to be less methylated then adult DNA. This difference has been proposed as a key factor in their ability to elicit inflammation (21, 24). We therefore characterised the methylation status of the cff-DNA acquired in our study. Although our data show a small percentage of unmethylated CpG motifs in cff-DNA, which was not detected in adult DNA when compared to E-coli DNA, it is plausible that this cff-DNA is not sufficiently hypomethylated to be pro-inflammatory, and to stimulate TLR9. Irrespective of this, neither our *in-vitro* or *in-vivo* studies support a simple model of cff-DNA acting as a pro-inflammatory agent in the induction of sPTB.

One limitation of our study is the caveat that the *in-vitro* and *in-vivo* models cannot represent the full complexity of human pathophysiology of sPTB. Placental DNA was chosen as a stimulus because cff-DNA (using a placental explant method) does not yield enough DNA for the *in-vivo* studies. Our *in-vivo* ultrasound guided intra-uterine model of sPTB was chosen because it is minimally invasive, allows for a more precisely targeted injection, and because LPS-injected animals have previously shown high rates of sPTB (26). We hypothesised that this model is more physiologically relevant than the intra-peritoneal injection of stimuli used in previous studies, which found that neither CpG nor mouse placental DNA induced adverse pregnancy outcomes in wildtype mice (27-30). This is the first study that examined the *in-vivo* effects of placental-derived mouse DNA on PTB / time to delivery. Our *in-vivo* study has a relatively small number of mice per group. However, there is a large difference in means between the placental DNA treated mice and the LPS treated mice. This leads to a more confident interpretation that intra-uterine placental DNA administration does not decrease time to delivery.

Our *in-vivo* findings conflict with findings from Scharfe and colleagues who found that human fetal DNA caused increased IL-6 production in PBMCs from pregnant women and caused fetal resorption in a murine model *in-vivo* (7). However, our findings are consistent with more recent literature, showing that various types of DNA (mouse adult and fetal and human fetal DNA) given through daily intra-peritoneal injections between 14 and 18 days of gestation, did not lead to higher fetal resorption in mice compared to control animals, whereas LPS injected animals showed 100% resorption (30). Furthermore, 3 separate studies showed no effects on fetal resorption in wildtype mice when given single intraperitoneal injections of the TLR9 ligand CpG ODN at 25-400 μg on days 10 or 14 of gestation (27-29).

The different findings in the above studies might be a consequence of the different methodologies used. Our *in-vivo* model used mouse placental DNA, as this is more comparable to the placental derived cff-DNA found in human maternal circulation and does not add the confounding factor of crossing species. In contrast the mouse model utilised by Scharfe and colleagues used human fetal DNA as the stimulus. Moreover, C57Bl/6 mice were used in our model and in another study that similarly showed no adverse pregnancy outcomes to various types of DNA (30), while Scharfe and colleagues used a BALB/c strain (7). The timing and frequency of injections also differs throughout studies; we administered a single injection at day 17 of gestation (because LPS injection at this time point leads to preterm delivery (26)), whereas Scharfe and colleagues gave a single injection between days 10 and 14 of gestation. Interestingly, it may be worth noting that despite their findings of fetal resorption, the systemic inflammation that is seen in studies using LPS-induced sPTB models (26) was not observed after administration of fetal DNA. It is possible that the human fetal DNA used could not induce sufficient inflammation to cause sPTB or that the fetal resorption observed was mediated by a pathway induced by injection between days 10 and 14 of gestation, but with a different outcome when stimuli were not introduced until day 17. Finally, the concentration of DNA used by Scharfe and colleagues, at which fetal resorption occurred (300 μg), must be considered highly supraphysiological, with 6 ng/ml being one of the highest concentration of cff-DNA measured in vivo during human pregnancy (13).

The mechanisms that underpin the generation of cff-DNA in pregnancy remain to be determined, as does the contribution of placental inflammation, often caused by infection in sPTB (4, 31), to this process. LPS has been shown to increase apoptosis rates in placental trophoblasts (32) and mediate cff-DNA release from mouse placental explants (20). However, in our placental explant model, using non-labouring term human placentas, we found no evidence that LPS-treated placental explants released higher amounts of cff-DNA compared to control samples, despite producing TNF-α in a dose dependent manner. This suggests that LPS-induced inflammation alone may not be sufficient to cause a clinically relevant increase in cff-DNA.

## Conclusion

Based on our findings we propose that cff-DNA alone is not pro-inflammatory, and is not sufficient to cause sPTB *in-vivo.* This suggests that the rise in cff-DNA measured in women who deliver preterm compared to women who deliver term, might be predominantly an effect of parturition and not a principal causative.

## Acknowledgements

This research was made possible by funding from Tommy’s.

Donald J Davidson was supported by a Medical Research Council Senior Non-clinical Fellowship (G1002046).

## Contribution of authors

Sara van Boeckel designed and conducted the experiments, interpreted the data and wrote the manuscript.

Heather MacPherson assisted in the conduction of *in-vivo* experiments, contributed to the experimental design, interpretation of data and writing of the manuscript.

DJD contributed to experimental design, interpretation of data and writing of the manuscript.

Jane E Norman contributed to the writing of the manuscript.

Sarah J Stock designed the experiments, contributed to the interpretation of data and writing of the manuscript.

